# Cognitive and affective neurodevelopment in youth exposed to deprivation and threat

**DOI:** 10.64898/2026.06.25.734615

**Authors:** Omid Kardan, Mike Angstadt, M. Fiona Molloy, Elisa M. Trucco, Mary M. Heitzeg, Katherine L. McCurry

## Abstract

**Background:** Early life adversity (ELA) is associated with notable negative consequences across development. Experiences of deprivation may affect neurocognitive development, while experiences of threat may alter emotion processing. Deprivation and threat may also differentially influence reward processing. However, unique consequences of deprivation and threat beyond low family resources are debated.

**Methods:** We employed an exposure vs. control data analytic approach to isolate deprivation and threat influences from socioeconomic resources. Adolescent Brain Cognitive Development (ABCD^®^) Study youth exposed to neither deprivation nor threat (N=2408-2962) were matched to youth exposed to deprivation-only (N=638-721), threat-only (N=198-232), or threat non-exclusively (threat+: N=382-464) based on family income, parental education, race/ethnicity, sex, and age. Multivariate analyses were used to distinguish each ELA group from their respective control groups in the *<u>neurocognitive domain</u>* (resting-state connectomic maturation, cognitive task performance, and cortical grey matter thickness at two timepoints) and in the *<u>neuroaffective domain</u>* (nucleus accumbens and caudate activation to reward anticipation and amygdala and insula activation to fearful faces).

**Results:** In the *<u>neurocognitive domain</u>*, similar latent variables (LVs) differentiated the deprivation and threat+ groups from their respective matched control groups. This LV corresponded to neurocognitive maturation, loading positively on cortical functional maturation and task performance, and negatively on cortical grey matter thickness. This LV was weaker in the deprivation and threat+ groups compared to controls. In the *<u>neuroaffective domain</u>*, no significant LVs were found.

**Conclusion:** Both threat and deprivation exposure during childhood may delay neurocognitive development in early adolescence beyond their co-occurrence with low socioeconomic resources.

## Introduction

Early life adversity (ELA) is associated with notable negative consequences across development including increased likelihood of drug use during adolescence (1), greater risk of developing a substance use disorder (SUD) in young adulthood (2), and increased risk of mental health problems across the lifespan (3,4). Because exposure occurs during a period of significant brain development and neuroplasticity (5), ELA may kick off a cascade of changes that predispose individuals to negative outcomes that accumulate across the lifespan (6). A deeper understanding of the impact of these experiences on neurodevelopment may help inform early intervention. However, experiences of ELA are heterogenous (e.g., neglect vs. abuse) and may impact different neurodevelopmental pathways (e.g., cognitive vs. affective) in distinct ways (7). Additionally, these experiences frequently take place within the context of other stressors such as limited socioeconomic resources (8–10), making it difficult to examine unique effects of these dimensions without large, well-characterized samples. Here, we utilize data from a large cohort of youth to investigate the impacts of different ELA dimensions—specifically deprivation and threat—on neurocognitive and neuroaffective development in early adolescence. We employ an exposure vs. matched data analytic approach to allow for a more specific isolation of deprivation and threat influences over those attributable to socioeconomic resources more generally.

The dimensional model of adversity and psychopathology suggests one meaningful way of parsing the heterogeneity in ELA is by considering dimensions underlying different ELA experiences with threat and deprivation representing two important dimensions (11,12). In this model, experiences of deprivation, characterized by lack of expected inputs or cognitive or social stimulation, may result in deficits in cognitive processing and altered frontoparietal structure and function (8). In contrast, experiences of threat, characterized as actual or potential harm or violence, may impact emotion processing, such as heightened vigilance towards potential hazards and exaggerated responsivity of limbic regions and salience networks (8). Reward processing may also be impacted by ELA (13), with deprivation and threat potentially exerting opposite influences on striatal activation (8). While there is a growing body of evidence supporting aspects of this model, few studies have examined the impact of these exposures on both cognitive and affective neurodevelopment within the same youth cohort.

Individuals with ELA frequently exhibit altered cognitive functioning (14,15) with accumulating evidence suggesting deprivation may have a more profound impact on cognition than threat (16,17). For example, in a cohort of US adults, those with court-documented records of childhood maltreatment had poorer cognitive performance compared to those without, and follow-up analyses showed this was significant in those with documented neglect but not in those with documented physical or sexual abuse (17). Similarly, a meta-analysis of studies focused on children and adolescents found impaired executive functioning was linked to both threat and deprivation exposure, but for working memory and inhibitory control, associations were stronger for deprivation than for threat (16). The neurodevelopmental changes that lead to these deficits are less well-understood. Across studies of adults who experienced ELA, diminished grey matter volume in the hippocampus, amygdala, and frontal regions (18), as well as reduced cortical thickness (19) have been consistently found. However, these patterns appear to be sensitive to age, with stronger effect sizes seen in studies with older samples (18,20), suggesting greater heterogeneity in studies of youth and pointing to the importance of understanding earlier developmental trajectories.

A second prominent pathway by which ELA has been hypothesized to contribute to heightened risk for negative outcomes is through its impact on affective processing (21,22). Individuals who have experienced ELA often exhibit alterations in both threat (8,23) and reward (13,24) processing. Across neuroimaging studies of emotional face processing, individuals exposed to childhood maltreatment demonstrate heightened responsivity in the amygdala, insula, parahippocampal and superior temporal gyri compared to those who have not experienced childhood maltreatment (23), with exaggerated amygdala and anterior insula reactivity also being seen in other tasks probing threat processing (8). Exposure to deprivation in childhood has generally been associated with diminished behavioral (25,26) and neural responsivity to rewards, particularly in the striatum (27–29). However, in individuals exposed to childhood threat, reward-related alterations have been less consistent, with some indicating decreased responsivity (30,31), while others have shown increased responses to reward (32,33) or no association (28).

Children living in households with low socioeconomic resources are at heightened risk for ELA (34,35); however, not all children from households with low socioeconomic resources experience deprivation or threat. Moreover, limited socioeconomic resources in childhood have also been linked to reduced executive functioning (36,37) and alterations of the developing brain (10,38–40). Thus, in previous work that did not account for demographic differences, differences attributed to threat or deprivation may instead be driven by group-level differences in socioeconomic resources. Importantly, some of these group differences may not be sufficiently accounted for via the addition of socioeconomic covariates in the regression models due to extreme propensity differences (see Methods).

To allow for a more specific isolation of deprivation and threat influences over those attributable to socioeconomic resources more generally, we utilized the large sample size in the Adolescent Brain Cognitive Development (ABCD) Study^®^ and identified youth exposed to neither deprivation nor threat who were demographically-matched with youth in the deprivation and threat groups. Matching was based on family income, parental education, race/ethnicity, sex, and age. These matched control groups were subsequently used in multivariate analyses investigating the neurocognitive and neuroaffective correlates of deprivation and threat. In the neurocognitive domain, we hypothesized that deprivation would be related to reduced neurocognitive development (reduced cognitive performance/delayed cortical maturation/reduced cortical thinning) above and beyond general socioeconomic resources (household income and parental education), while threat, in isolation, may not be as strongly related to neurocognitive development. In the neuroaffective domain, we hypothesized that threat would be related to altered amygdala and insula responsivity to fearful faces, while deprivation would be associated with diminished striatal responsiveness to reward anticipation.

## Methods

### Participants

We utilized data from the ABCD Study (41,42), which is an ongoing longitudinal study of 11,867 children across 21 sites in the U.S. Participants were enrolled in the study between 9–10.9 years of age (Y0), and our analyses additionally utilized their two-year follow-up (Y2; 11-12.9 years) data. In analyses examining neurocognitive development, a total of 3645 participants were included with neuroimaging and neurocognitive measures at both Y0 and Y2 (see resting-state fMRI exclusions section). Of these, 116 who did not respond to one or more items assessing threat/deprivation were excluded. In analyses investigating neuroaffective development, 4344 participants with neuroaffective measures at both Y0 and Y2 were assessed (see task-fMRI exclusions section). Of these, 124 who did not respond to one or more of the threat/deprivation items were excluded.

For each analysis, matched control groups were drawn from the pool of participants with no exposure to deprivation or threat (N = 2408 in neurocognitive and N = 2962 in neuroaffective analysis). **Table 1** shows demographic characteristics of the groups of participants reporting deprivation exclusively and their matched control groups in each analysis. **Table 2** shows these for participants exclusively exposed to threat and their matched control groups. Due to low occurrence of threat-only exposure, we also assessed a non-exclusive threat group (threat+ which includes participants who experienced threat regardless of whether they also experienced deprivation or not). **Table 3** shows demographic characteristics of the threat+ groups for each analysis and the matched control groups. Details about the matching process are reported in the **Supplementary section ‘Demographic matching’**.

**Table 1.** Demographic break-down of the participants within the deprivation groups and their matched control groups for the neurocognitive (*top*) and the neuroaffective (*bottom*) analyses. *Notes*: Mean household income was approximated by interpolating mean of the midpoints of income brackets. Min and max income are shown in brackets. Education is based on the highest parental years of education. Bold values show Cohen’s d > 0.2 prior to matching.

| **Neurocognitive Analysis** | Deprivation group<br>(N = 638) | Control group<br>matched with<br>Deprivation<br>group<br>(N = 534) | No deprivation<br>or threat pool of<br>participants<br>(N = 2408) | Cohen's d<br>(Deprivation<br>vs. Control) | Cohen's d<br>(Deprivation<br>vs. Pool) |
| --- | --- | --- | --- | --- | --- |
| **Parental Education** | 16.7 yrs<br>(SD = 2.5) | 16.9 yrs<br>(SD = 2.4) | 17.9 yrs<br>(SD = 2.1) | -0.07 | **-0.36** |
| **Household Income** | \$58.7K [5K-200K] | \$66.7K [5K-200K] | \$101K [5K-200K] | -0.12 | **-0.51** |
| **White** | 283 (44.4%) | 266 (49.8%) | 1654 (68.7%) | -0.08 | **-0.36** |
| **Hispanic** | 163 (25.5%) | 121 (22.7%) | 352 (14.6%) | 0.05 | **0.20** |
| **Black** | 115 (18.0%) | 83 (15.5%) | 164 (6.8%) | 0.05 | **0.24** |
| Asian/Other | 77 (12.1%) | 64 (12.0%) | 238 (9.9%) | 0.00 | 0.05 |
| Female | 314 (49.2%) | 257 (48.1%) | 1202 (49.9%) | 0.01 | -0.01 |
| Age at Y0 | 9.90 yrs<br>(SD = 0.63) | 9.92 yrs<br>(SD = 0.63) | 9.96 yrs<br>(SD = 0.62) | -0.01 | -0.06 |
| Age at Y2 | 11.94 yrs<br>(SD = 0.67) | 11.97yrs<br>(SD = 0.67) | 12.01 yrs<br>(SD = 0.65) | -0.03 | -0.08 |
| **Neuroaffective Analysis** | Deprivation group<br>(N = 721) | Control group<br>matched with<br>Deprivation<br>group<br>(N = 603) | No deprivation<br>or threat pool of<br>participants<br>(N = 2962) | Cohen's d<br>(Deprivation<br>vs. Control) | Cohen's d<br>(Deprivation<br>vs. Pool) |
| **Parental Education** | 16.8 yrs<br>(SD = 2.4) | 17.0 yrs<br>(SD = 2.4) | 18.0 yrs<br>(SD = 2.1) | -0.08 | **-0.38** |
| **Household Income** | \$58.3K [5K-200K] | \$66.6K [5K-200K] | \$107K [5K-200K] | -0.12 | **-0.55** |
| **White** | 328 (45.5%) | 300 (49.8%) | 2208 (68.5%) | -0.06 | **-0.34** |
| Hispanic | 175 (24.3%) | 152 (23.6%) | 430 (14.5%) | 0.01 | 0.18 |
| **Black** | 131 (18.2%) | 91 (15.1%) | 186 (6.3%) | 0.06 | **0.26** |
| Asian/Other | 87 (12.1%) | 70 (11.6%) | 318 (10.7%) | 0.01 | 0.03 |
| Female | 329 (45.6%) | 260 (43.1%) | 1386 (46.8%) | -0.04 | 0.02 |
| Age at Y0 | 9.92 yrs<br>(SD = 0.63) | 9.89 yrs<br>(SD = 0.63) | 9.95 yrs<br>(SD = 0.62) | 0.03 | -0.04 |
| Age at Y2 | 11.97 yrs<br>(SD = 0.66) | 11.96 yrs<br>(SD = 0.67) | 12.03 yrs<br>(SD = 0.65) | 0.00 | -0.06 |

**Table 2.**
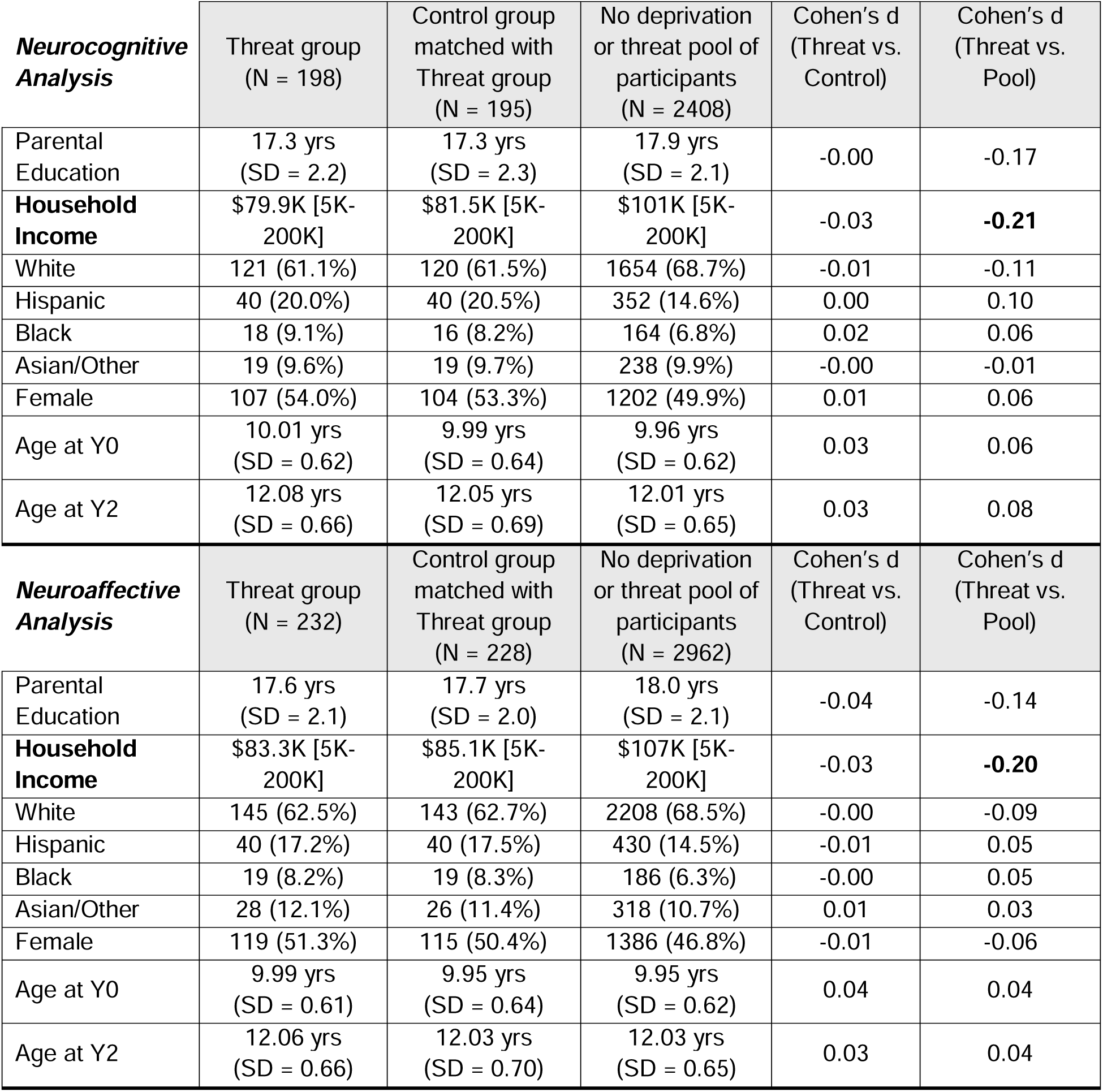
Demographic break-down of the participants within the threat groups and their matched control groups for the neurocognitive (*top*) and the neuroaffective (*bottom*) analyses. *Notes*: Mean household income was approximated by interpolating mean of the midpoints of income brackets. Min and max income are shown in brackets. Education is based on the highest parental years of education. Bold values show Cohen’s d > 0.2 prior to matching.

| **Neurocognitive Analysis** | Threat group<br>(N = 198) | Control group<br>matched with<br>Threat group<br>(N = 195) | No deprivation<br>or threat pool of<br>participants<br>(N = 2408) | Cohen's d<br>(Threat vs.<br>Control) | Cohen's d<br>(Threat vs.<br>Pool) |
| --- | --- | --- | --- | --- | --- |
| Parental Education | 17.3 yrs<br>(SD = 2.2) | 17.3 yrs<br>(SD = 2.3) | 17.9 yrs<br>(SD = 2.1) | -0.00 | -0.17 |
| **Household Income** | \$79.9K [5K-200K] | \$81.5K [5K-200K] | \$101K [5K-200K] | -0.03 | **-0.21** |
| White | 121 (61.1%) | 120 (61.5%) | 1654 (68.7%) | -0.01 | -0.11 |
| Hispanic | 40 (20.0%) | 40 (20.5%) | 352 (14.6%) | 0.00 | 0.10 |
| Black | 18 (9.1%) | 16 (8.2%) | 164 (6.8%) | 0.02 | 0.06 |
| Asian/Other | 19 (9.6%) | 19 (9.7%) | 238 (9.9%) | -0.00 | -0.01 |
| Female | 107 (54.0%) | 104 (53.3%) | 1202 (49.9%) | 0.01 | 0.06 |
| Age at Y0 | 10.01 yrs<br>(SD = 0.62) | 9.99 yrs<br>(SD = 0.64) | 9.96 yrs<br>(SD = 0.62) | 0.03 | 0.06 |
| Age at Y2 | 12.08 yrs<br>(SD = 0.66) | 12.05 yrs<br>(SD = 0.69) | 12.01 yrs<br>(SD = 0.65) | 0.03 | 0.08 |
| **Neuroaffective Analysis** | Threat group<br>(N = 232) | Control group<br>matched with<br>Threat group<br>(N = 228) | No deprivation<br>or threat pool of<br>participants<br>(N = 2962) | Cohen's d<br>(Threat vs.<br>Control) | Cohen's d<br>(Threat vs.<br>Pool) |
| Parental Education | 17.6 yrs<br>(SD = 2.1) | 17.7 yrs<br>(SD = 2.0) | 18.0 yrs<br>(SD = 2.1) | -0.04 | -0.14 |
| **Household Income** | \$83.3K [5K-200K] | \$85.1K [5K-200K] | \$107K [5K-200K] | -0.03 | **-0.20** |
| White | 145 (62.5%) | 143 (62.7%) | 2208 (68.5%) | -0.00 | -0.09 |
| Hispanic | 40 (17.2%) | 40 (17.5%) | 430 (14.5%) | -0.01 | 0.05 |
| Black | 19 (8.2%) | 19 (8.3%) | 186 (6.3%) | -0.00 | 0.05 |
| Asian/Other | 28 (12.1%) | 26 (11.4%) | 318 (10.7%) | 0.01 | 0.03 |
| Female | 119 (51.3%) | 115 (50.4%) | 1386 (46.8%) | -0.01 | -0.06 |
| Age at Y0 | 9.99 yrs<br>(SD = 0.61) | 9.95 yrs<br>(SD = 0.64) | 9.95 yrs<br>(SD = 0.62) | 0.04 | 0.04 |
| Age at Y2 | 12.06 yrs<br>(SD = 0.66) | 12.03 yrs<br>(SD = 0.70) | 12.03 yrs<br>(SD = 0.65) | 0.03 | 0.04 |

**Table 3.** Demographic break-down of the participants within the threat+ groups and their matched control groups for the neurocognitive (*top*) and the neuroaffective (*bottom*) analyses. *Notes*: Mean household income was approximated by interpolating mean of the midpoints of income brackets. Min and max income are shown in brackets. Education is based on the highest parental years of education. Bold values show Cohen’s d > 0.2 prior to matching.

| **Neurocognitive** | Threat+ group<br>(N = 382) | Control group<br>matched with<br>Threat+ group<br>(N = 346) | No deprivation<br>or threat pool of<br>participants<br>(N = 2408) | Cohen's d<br>(Threat+<br>vs. Control) | Cohen's d<br>(Threat+ vs.<br>Pool) |
| --- | --- | --- | --- | --- | --- |
| **Parental Education** | 16.9 yrs<br>(SD = 2.1) | 17.0 yrs<br>(SD = 2.1) | 17.9 yrs<br>(SD = 2.1) | -0.04 | **-0.32** |
| **Household Income** | \$62.0K [5K-200K] | \$67.6K [5K-200K] | \$101K [5K-200K] | -0.08 | **-0.48** |
| **White** | 203 (53.2%) | 192 (55.5%) | 1654 (68.7%) | -0.03 | **-0.23** |
| Hispanic | 74 (19.4%) | 67 (19.4%) | 352 (14.6%) | 0.00 | 0.09 |
| Black | 58 (15.2%) | 45 (13.0%) | 164 (6.8%) | 0.04 | 0.19 |
| Asian/Other | 47 (12.3%) | 42 (12.1%) | 238 (9.9%) | 0.00 | 0.05 |
| Female | 196 (51.3%) | 176 (50.9%) | 1202 (49.9%) | -0.00 | -0.02 |
| Age at Y0 | 9.98 yrs<br>(SD = 0.63) | 10.00 yrs<br>(SD = 0.63) | 9.96 yrs<br>(SD = 0.62) | -0.02 | 0.02 |
| Age at Y2 | 12.03 yrs<br>(SD = 0.66) | 12.04 yrs<br>(SD = 0.68) | 12.01 yrs<br>(SD = 0.65) | -0.01 | 0.02 |
| **Neuroaffective** | Threat+ group<br>(N = 464) | Control group<br>matched with<br>Threat+ group<br>(N = 423) | No deprivation<br>or threat pool of<br>participants<br>(N = 2962) | Cohen's d<br>(Threat+<br>vs. Control) | Cohen's d<br>(Threat+ vs.<br>Pool) |
| **Parental Education** | 16.9 yrs<br>(SD = 2.1) | 17.1 yrs<br>(SD = 2.2) | 18.0 yrs<br>(SD = 2.1) | -0.06 | **-0.38** |
| **Household Income** | \$60.9K [5K-200K] | \$65.9K [5K-200K] | \$107K [5K-200K] | -0.08 | **-0.53** |
| **White** | 240 (51.7%) | 222 (52.5%) | 2208 (68.5%) | -0.01 | **-0.25** |
| Hispanic | 87 (18.8%) | 84 (19.9%) | 430 (14.5%) | -0.02 | 0.08 |
| Black | 66 (14.2%) | 58 (13.7%) | 186 (6.3%) | 0.01 | 0.19 |
| Asian/Other | 71 (15.3%) | 59 (14.0%) | 318 (10.7%) | 0.03 | 0.10 |
| Female | 228 (49.1%) | 206 (48.7%) | 1386 (46.8%) | -0.00 | -0.03 |
| Age at Y0 | 9.97 yrs<br>(SD = 0.61) | 9.98 yrs<br>(SD = 0.62) | 9.95 yrs<br>(SD = 0.62) | -0.00 | 0.02 |
| Age at Y2 | 12.02 yrs<br>(SD = 0.65) | 12.06 yrs<br>(SD = 0.66) | 12.03 yrs<br>(SD = 0.65) | -0.04 | 0.00 |

The deprivation group, threat-only group, and threat+ group can be seen in the Venn diagram in **Figure 1** (deprivation is pink and outside the intersection, threat-only is light orange outside the intersection and threat+ is all of the light and dark orange circle).

**Figure 1.**
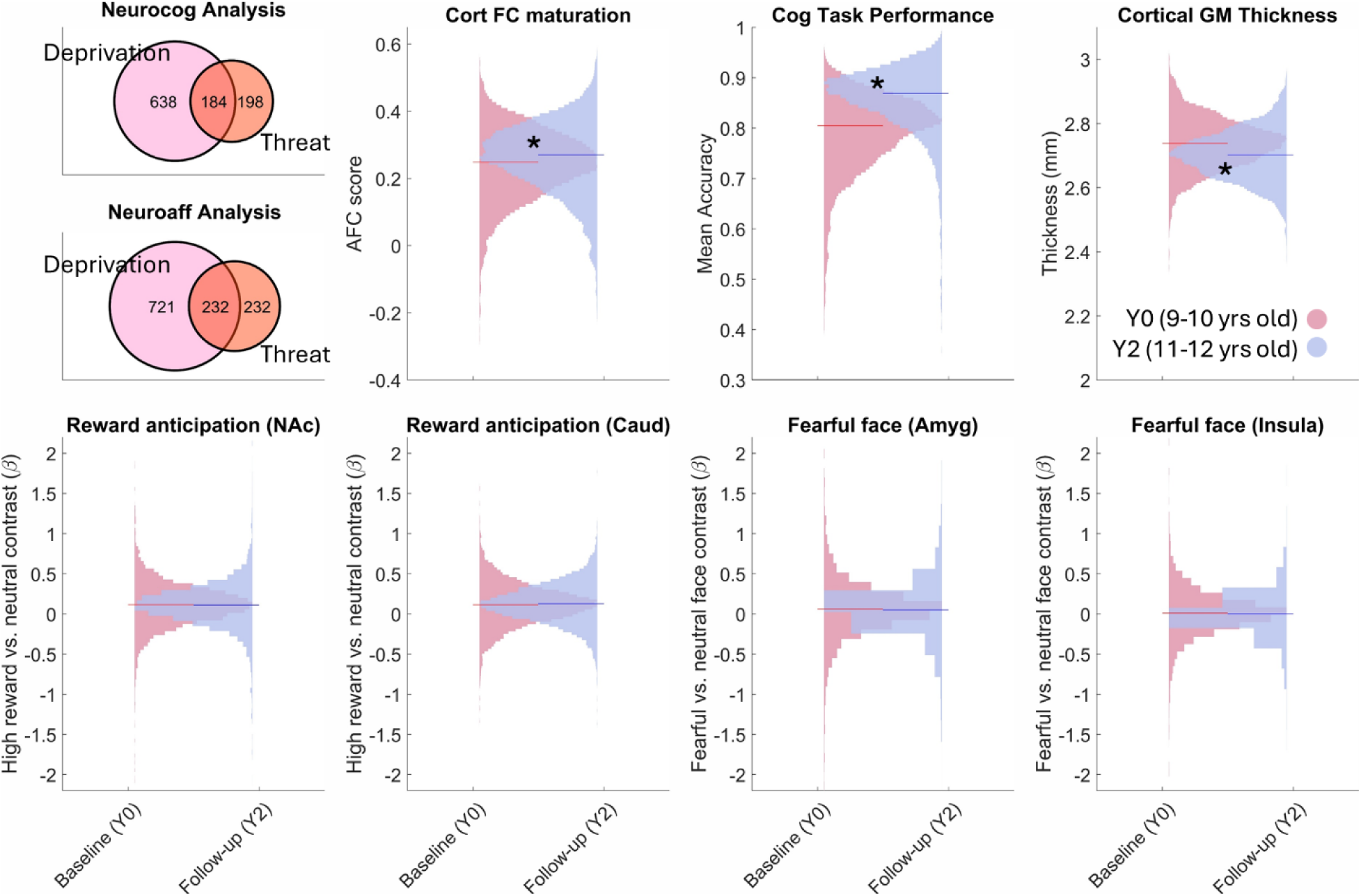
**Top-left panel:** Venn diagram of deprivation and threat exposure (separately for the neurocognitive and neuroaffective analysis). The deprivation group is shown in pink and outside the intersection, the threat-only group is shown in light orange outside the intersection, and the threat+ group is composed of all of the light and dark orange circle. **Other panels:** The distributions of the neurocognitive (*top row*) and neuroaffective (*bottom row*) measures in the full pool of participants of the current study across the baseline (*red*) and follow-up (*blue*) years. * indicates a significant (p < .01 based on paired t-test) increase or decrease in the measure between the two waves. *Abbreviations*: AFC = anchored functional connectivity; Cort FC = cortical functional connectivity; Cog = cognitive; GM = gray matter; NAc = nucleus accumbens; Caud = caudate; Amyg = Amygdala.

### Deprivation and threat exposure variables

We utilized a multi-measure, multi-informant approach to calculating experiences of deprivation and threat that was informed by ELA theory (e.g., (43)) and prior topical work with the ABCD Study sample (e.g., (44,45)). Individuals with presence of one or more of these indicators at Y0, Y1, or Y2 were classified as positive cases (i.e., deprivation exposure or threat exposure). Briefly, five youth-reported items from the Acceptance scale of the Children’s Report of Parental Behavior Inventory (46,47) were used as a proxy measure of <u>emotional deprivation</u>, two youth-reported items from the Parental Monitoring Questionnaire (48) were used as a proxy of <u>supervisory deprivation</u>, and four parent-reported items about needing food but being unable to afford it, being evicted, having gas or electricity turned off, and needing to see a doctor but not able to afford it were used as proxies of <u>physical deprivation</u>. While the physical deprivation items are related to household income, they reflect direct experiences of deprivation and do not occur in all low-income households. To assess <u>threat exposure</u>, ten parent-reported items from the Kiddie Schedule for Affective Disorders and Schizophrenia for School-Aged Children (KSADS-COMP) were assessed including items measuring verbal threats of death, physical abuse, sexual abuse, and witnessing domestic or community violence. More details can be found in the **Supplementary section ‘Adversity exposure variables.’**

### Resting-state and task fMRI data

Resting-state **(rsfMRI)**, Monetary incentive delay task **(MID)** (49), and Emotional n-back task **(EN-back)** (41) fMRI at Y0 and Y2 were used to construct measures of cortical network maturation, reward processing, and threat processing, respectively (see neurocognitive and neuroaffective variables section). More details about fMRI data processing and the tasks can be found in the **Supplementary section ‘Resting-state and task fMRI data.’**

### Neurocognitive variables

#### Cortical functional connectomic maturation

The maturity of the participant’s cortical networks was calculated as their connectome’s proximity to adult and distance from baby networks. Specifically, the Anchored Functional Connectivity maturation (**AFC**) score (50) was calculated for the resting-state functional connectivity matrix of each participant at each timepoint (Y0 and Y2). As the boundaries of brain functional networks during adolescence are in transition from early-life to adulthood, AFC captures normative functional maturation in youth by comparing the distance of the connectome to an early-life set of networks (51) vs. its proximity to an adult set of networks (52). Due to its two anchoring points, AFC is significantly better than distance from adult networks alone in capturing age and aging in youth (50). Importantly, cognitive performance across and within youth are associated with AFC differences across and changes in AFC within youth, respectively (50).

Specifically, AFC was calculated as follows for each participant at Y0 as well as at Y2:

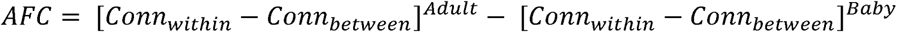

Above, [Conn_within –_ Conn_between_]^Adult^ is the average connectivity within networks minus the average connectivity between networks specified by (52), while [Conn_within –_ Conn_between_]^Baby^ is the average connectivity within networks minus the average connectivity between networks specified by (51). These sets of networks and their anatomical maps, their validity for youth rsfMRI data, and examples of AFC comparisons between two participants in the ABCD Study sample are shown in our previous works (50,53,54).

#### Cortical thickness

The average thickness in mm over all cortical parcels of both hemispheres based on the Desikan-Killiany atlas (55) was calculated for Y0 and Y2 of each participant based on the generated variable ‘smri_thick_cdk_mean’ by the ABCD Data Analysis, Informatics & Resource Center (DAIRC). Mean cortical thickness normatively starts to decline in close to linear fashion after toddlerhood during development (56). Our previous work using the ABCD Study sample reported a normative increase in AFC and decrease in cortical thickness from Y0 to Y2 (i.e., from 9-10 years of age to 11-12 years of age) (53), and see **Figure 1** for the participants in the current study.

#### Cognitive task performance

Cognitive performance was quantified as average performance in the EN-back task (41), and the five out of seven NIH-Toolbox tasks (57) that were available longitudinally at both Y0 and Y2: the Picture Vocabulary task, Flanker inhibitory control and attention task, Pattern Comparison processing speed task, Picture sequence memory task, and Oral reading recognition task. This composite shows a robust development from Y0 to Y2 (e.g., see (53) or **Figure 1**) and captures a wide range of developing cognitive processes in youth including sustained attention and working memory (58).

### Neuroaffective variables

#### ROI-specific activation in response to fearful faces

Negative emotional reactivity was estimated based on the contrast beta weights of fearful face vs. neutral face trials in the EN-back task in amygdala using Freesurfer segmentation and insula based on the Desikan-Killiany cortical atlas. These variables are provided by the DAIRC as part of the mri_y_tfmr_nback_ngfvntf_aseg and mri_y_tfmr_nback_ngfvntf_dsk data tables. The distributions of these values in the current study can be seen in **Figure 1**.

#### ROI-specific activation in anticipation of reward

Reward anticipation in nucleus accumbens and caudate (based on Freesurfer automatic segmentation in both right and left hemispheres) were estimated as the contrast beta weights for anticipation of large reward versus neutral trials of the MID task. These variables are provided by the DAIRC as part of the mri_y_tfmr_mid_alrvn_aseg data table.

### Covariates

In addition to the matching variables (age, sex, race/ethnicity, family income, and parental education), results were adjusted for head motion (mean frame displacement in the EN-back and MID fMRI runs for the neuroaffective analyses and mean frame displacement for the rsfMRI runs for the neurocognitive analyses), site, scanner, number of included rsfMRI runs (only for neurocognitive analyses), neighborhood socioeconomic disadvantage as indicated by the Area Deprivation Index (59), and participant’s pubertal status (the average of youth and parental reports on the Pubertal Developmental Scale categorized sum score; 1: pre-puberty, 2: early puberty, 3: mid puberty, 4: late puberty, and 5: post puberty).

Given the potential association between early life adversities and neurodevelopmental and cognitive correlates of psychopathology (60,61), in sensitivity analyses we tested if additionally including the general factor of psychopathology as a covariate influenced the results. The general factor of psychopathology was quantified using a bi-factor model fit to parent-rated Child Behavior Checklist items as described in (62).

### Patial Least Squares (PLS) analyses

In the neurocognitive analyses, we utilized a mean-centered task PLS multivariate analysis approach (63,64) to find the linear combinations of neurocognitive variables (AFC score, cortical thickness, cognitive task performance) and groups-by-time contrasts (adversity vs. matched control at Y0 and Y2) that would be maximally correlated. More details of this approach can be found in (65). Briefly, singular value decomposition was applied to the covariance between the neurocognitive variables and the group-by-time contrasts. Each of the identified combinations of weights is orthogonal to the others and will be referred to as a Latent Variable (LV), with the primary LV explaining the highest amount of covariance between the two sets (i.e. neurocognitive set and group-by-time set).

A permutation test was used to test the significance of each LV, where singular value decomposition was applied to 5000 null covariance matrices (i.e., with permuted neurocognitive variables). The singular value of the calculated LV was compared to the distribution of the null singular values to determine its significance. The partial correlations of the LV with the original neurocognitive variables and dummy variables coding the group-by-time contrasts were used to turn the weights into more interpretable loadings while also adjusting them for the covariates. Confidence intervals were built around these adjusted loadings via bootstrapping the sample 5000 times. We only interpreted LVs that passed both the permutation test (p < .05) and the bootstrap test (adjusted LV confidence intervals did not contain zero for at least one variable on each side).

The neuroaffective PLS was identical to the neurocognitive PLS but used the neuroaffective variables (anticipatory reward activation in NAc, anticipatory reward activation in caudate, fearful face processing in amygdala, fearful face processing in insula) instead of the neurocognitive variables.

## Results

### 1. Neurocognitive and neuroaffective development in deprivation exposure

The primary LV distinguishing the deprivation group from their matched control group over the two timepoints was significant in the <u>neurocognitive analysis</u> (permutation *p* < .001). This LV’s loadings on the neurocognitive measures are shown in the left panel of **Figure 2A**, while its adjusted loadings on the two groups (deprivation vs. matched control) across the two timepoints (9-10 [Y0] and 11-12 years of age [Y2]) are shown in the right panel of **Figure 2A**. Overall (i.e., averaged over timepoints), this LV indicated a group difference between the deprivation and matched control groups beyond socioeconomic/demographic factors such that cortical connectomic maturation (AFC) and cognitive task performance are higher and cortical thickness is lower in the control group. This group difference was small but statistically significant (difference between the two groups in the LV loading averaged over time: Δr_partial_ = .07; bootstrap *p* <.001), suggesting a delay-like pattern in neurocognitive maturation in the deprivation group compared to their matched peers (e.g., see normative trajectory for each measure in **Figure 1**). From Y0 to Y2, there was no significant difference in the amount of growth in this neurocognitive LV between the two groups (bootstrap *p* = .584 for the group-by-time interaction). Therefore, we did not find preliminary evidence for a widening or shrinking gap in neurocognitive maturation between the two groups within early adolescence.

**Figure 2.**
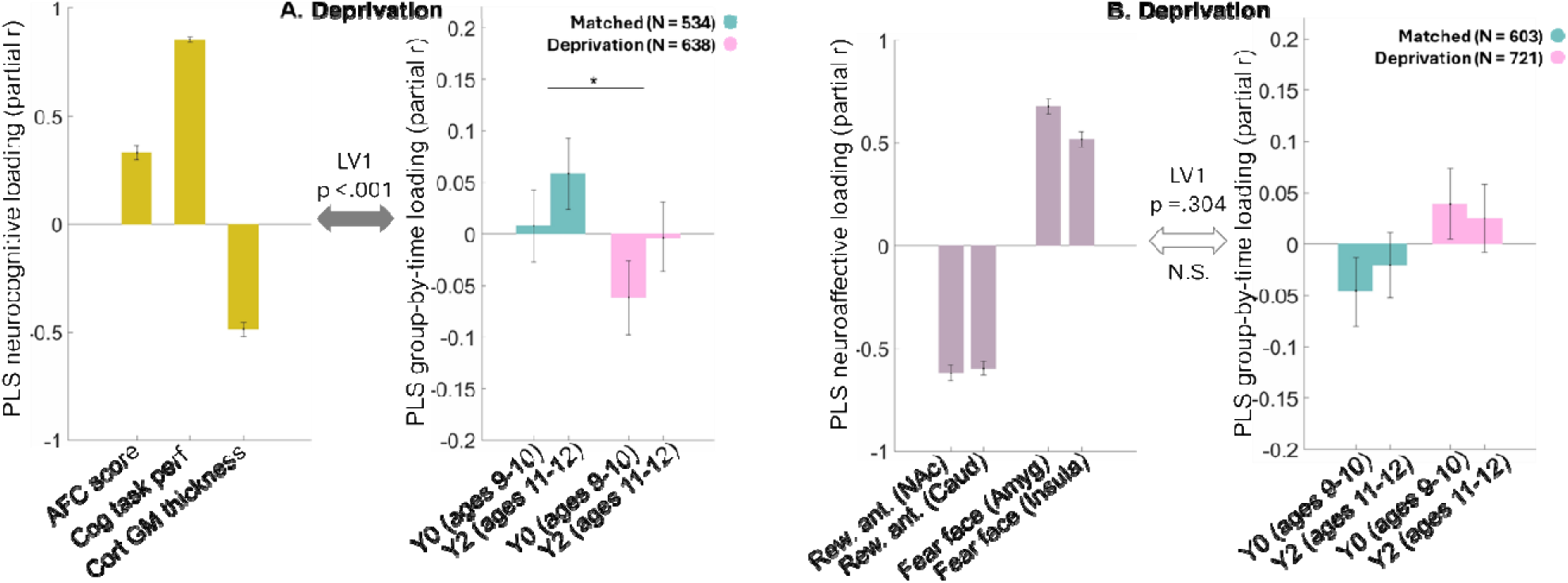
Deprivation vs. matched control analyses results. **A:** The y-axes show partial correlation of the PLS primary LV (adjusted for all covariates) with the neurocognitive measures (*left panel*) or group-by-time contrasts (*right panel*). Filled double-sided arrow indicates the LV is significant based on permutations. * within right panels indicates a significant group difference adjusted for all covariates. **B:** The y-axes show partial correlation of the LV (adjusted for all covariates) with the neuroaffective measures (*left panel*) or group-by-time contrasts (*right panel*). Empty double-sided arrow with N.S. indicates a non-significant LV (group contrasts are only evaluated if the LV is significant).

These analyses were adjusted for participants’ head motion, scanner, site, number of fMRI runs, family income, parental education, pubertal status at Y0 and Y2, age at Y0 and Y2, biological sex, race/ethnicity, and economic deprivation of the neighborhood (ADI). Excluding race/ethnicity from the covariates did not change the results (Δr_partial_ = .07; bootstrap *p* <.001). Adding a general factor of psychopathology as a covariate did not change the results (Δr_partial_ = .06; bootstrap *p* <.001).

In the <u>neuroaffective PLS analysis</u> (**Figure 2B**), the primary LV was not significant (permutation *p* = .075, N.S.). The trend of the LV was however consistent with hypo-activation in anticipation of reward and hyper-activation when processing fearful faces (i.e., lower anticipatory reward activation in NAc/Caudate and higher activation of Amygdala/Insula to fearful faces was found in the deprivation group compared to their matched control group).

### 2. Neurocognitive and neuroaffective development in threat exposure

<u>In the neurocognitive domain</u>, the analysis involving threat-only participants and their matched control group resulted in a primary LV that passed the permutation test (p < .001), but its loadings on the group-by-time contrast adjusted for covariates were unstable (all bootstrap confidence intervals cross zero; **Figure 3A**). Due to the small sample size of the threat-only group, we repeated the analysis with non-exclusive threat exposure (threat+) which roughly doubled the sample size for the analysis. Exposure to deprivation was added as a covariate in the threat+ analysis because the goal was to increase the sample size for threat exposure while accounting for deprivation. There is no overlap between the threat+ sample and the deprivation sample from the deprivation analyses reported above. As shown in **Figure 3B**, there was a delay-like pattern in neurocognitive development for the threat+ group compared to their matched control group such that higher AFC score, thinner cortical gray matter, and higher cognitive task performance loaded more strongly on the matched control group (Δr_partial_ = .09; bootstrap *p* <.001 for the difference between the two groups averaged over the two timepoints). Excluding race/ethnicity from the covariates did not change the results (Δr_partial_ = .09; bootstrap *p* <.001). Adding a general factor of psychopathology as a covariate did not change the results (Δr_partial_ = .08; bootstrap *p* <.001).

**Figure 3.**
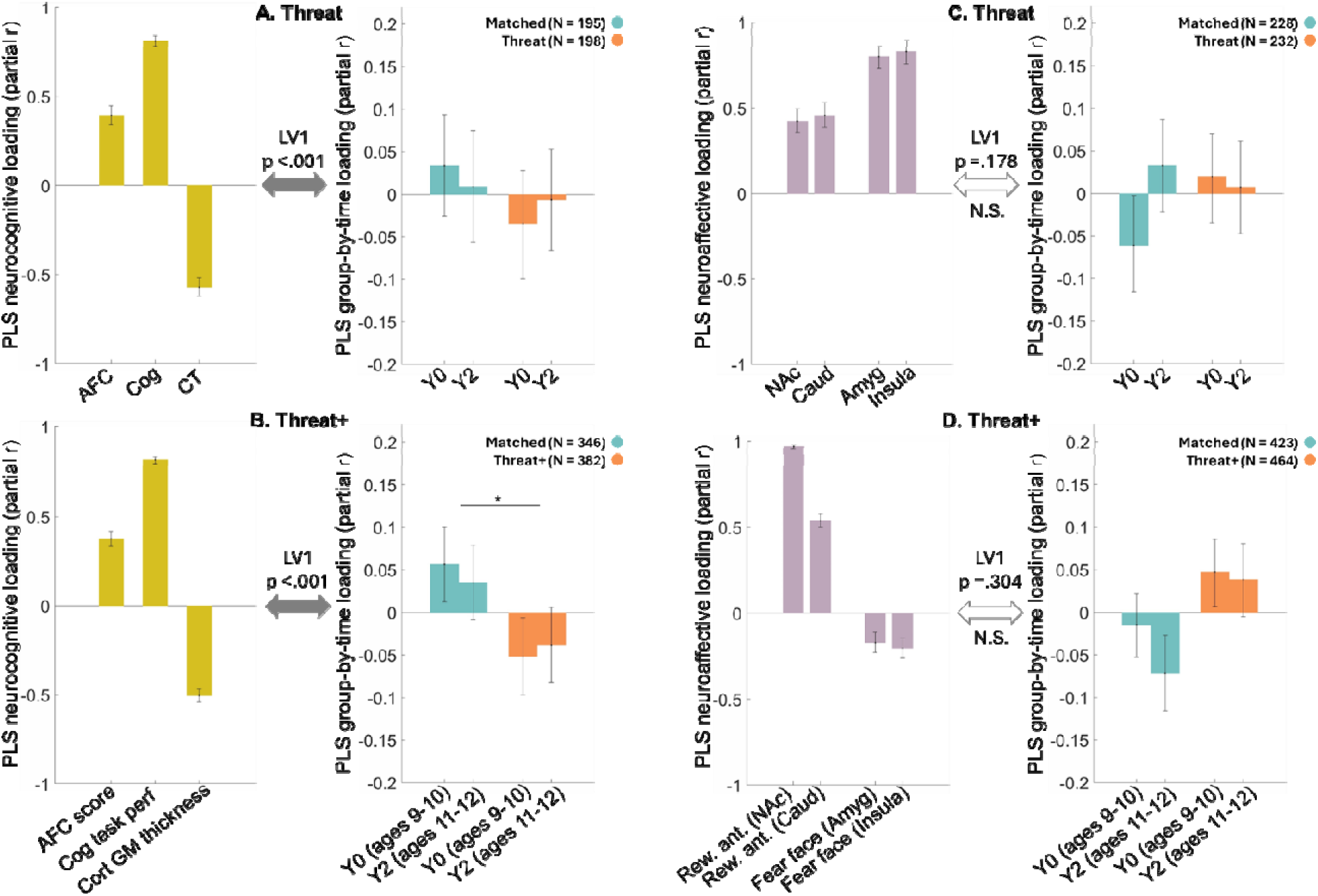
Threat and threat+ vs. matched control analyses results. **A:** The y-axes show partial correlations of the PLS primary LV (adjusted for all covariates) with the neurocognitive measures (left panel) or group-by-time contrasts (right panel) for the threat-only group vs. their matched control group. **B:** The y-axes show partial correlation of the LV adjusted for deprivation in addition to all covariates with the neurocognitive measures (left panel) or group-by-time contrasts (right panel) for the threat+ group vs. their matched control group. *within right panels indicates a significant group difference adjusted for all covariates and deprivation. **C:** The y-axes show partial correlation of the PLS primary LV (adjusted for all covariates) with the reward and threat processing neural measures (left panel) or group-by-time contrasts (right panel) for the threat-only group vs. their matched control group. **D:** The y-axes show partial correlation of the LV adjusted for deprivation in addition to all covariates with the reward and threat processing neural measures (left panel) or group-by-time contrasts (right panel) for the threat+ group vs. their matched control group. Empty double-sided arrows with N.S. indicate non-significant LVs which prevented us from evaluating the group contrasts.

In the neuroaffective domain, we found no significant or trending LVs in the PLS analyses for the threat-only (permutation *p* = .178, N.S.) or threat+ (permutation *p* = .304, N.S.) groups with regards to the reward anticipation and fearful face processing compared to their matched control groups (**Figure 3C** and **D**).

## Discussion

In this study, we identified a neurocognitive maturation profile that distinguished youth who experienced deprivation from matched peers without ELA exposure. Youth exposed to childhood deprivation showed poorer cognitive performance, greater cortical gray matter thickness, and less mature cortical functional connectivity than their socioeconomically matched peers without deprivation, a pattern consistent with delayed neurocognitive maturation. A similar neurocognitive profile distinguished youth exposed to threat, including those with co-occurring deprivation, from matched unexposed youth, even when accounting for deprivation. In contrast, neuroaffective processing in key reward- and threat-related regions did not distinguish youth exposed to threat from matched unexposed youth, nor did it distinguish youth exposed to deprivation from matched unexposed peers. Taken together, these findings suggest both threat and deprivation exposure during childhood are associated with delay-like patterns of neurocognitive development in early adolescence while neuroaffective processing in key limbic regions may not be robustly associated with threat or deprivation exposure during this developmental period.

Our finding of a delay-like pattern of neurocognitive maturation in deprivation exposed youth is broadly consistent with theories proposing limited inputs in childhood across a range of domains such as material, cognitive, and social, may alter neurodevelopment (5,66) as well as research demonstrating the impacts of deprivation on cognitive functioning (16,17) and structural and functional brain development (67–70). Notably, our work extends these findings by demonstrating that deprivation-related differences in neurocognitive maturation are evident above and beyond the impact of limited socioeconomic resources. Additionally, our measure of brain functional networks maturity does not assume that the adult canonical networks well-capture the brain functional organization at 9-12 years of age, which is a limitation of multiple studies assessing functional connectivity at the network level in youth samples. Recent work examining diffusion tensor imaging and resting-state functional connectivity data from the ABCD Study also found evidence of “younger-looking” brains associated with ELA experiences reflective of emotional neglect (71). Here, our maturation measures were theory-driven rather than data-driven, directly linked to cognitive performance development, and more interpretable than the learned features within brain-age models.

Within the context of threat, ELA has often been theorized to lead to accelerated neural development that may be adaptive to address short-term needs but suboptimal for later life functioning (66). Our threat-related neurocognitive findings do not align with this theorized threat-related accelerated development and instead suggest that early threat exposure may be associated with a delay-like pattern of neurocognitive development similar to that seen with deprivation exposure. The similar impact of deprivation and threat on structural brain metrics is supported by work demonstrating ELA-associated structural changes in adults at midlife were largely similar across exposures to threat or deprivation and not driven primarily by childhood socioeconomic resources (19). However, in this work, ELA was associated with thinner cortices at midlife rather than associated with thicker cortices as we found here, pointing to the importance of understanding how ELA impacts trajectories across the lifespan. Similarly, a meta-analysis of structural alterations associated with childhood maltreatment (20) found that age moderated the strength of regional differences, such that older adults showed greater reductions in cortical thickness and gray matter volume, suggesting that discrepancies between our threat-related finding and prior work could also be related to the age of the participants or the timing of adversity exposure (5).

Contrary to our expectation and to prior work demonstrating ELA-associated differences in affective processing (23,24), we did not find evidence that youth exposed to threat exhibited distinct patterns of neuroaffective processing in key brain regions from their matched unexposed peers. These null results are congruent with recent work that failed to find significant effects of threat exposure on neural correlates of emotion processing (72). Developmental differences between our study and prior work may be one reason for differences in findings; a recent meta-analysis of neural alterations associated with adversity exposure failed to find consistent patterns of altered neural response across studies focused on children or adolescents while replicating findings from prior work when examining studies focused on adults (73). Animal models of ELA suggest that there may be an incubation period following exposure in which behavioral and/or neural impacts are not yet fully evident (74). Another explanation for discrepancies may be that affective processing differences attributed to ELA may have been better explained by other factors such as socioeconomic resources. A strength of this study is the use of matched comparison groups allows for inferences to be made about the effects of ELA above and beyond these other potentially confounding factors. Finally, task-related differences may partially explain inconsistencies between some prior work and our null finding. Several of the previous studies that reported ELA or threat-related differences in amygdala responsivity used passive viewing tasks, which are more likely to elicit amygdala responses than other types of emotional tasks (75). In comparison, in this study, BOLD responses to emotional stimuli were measured within the context of an n-back task, reflective of implicit emotion regulation, which may have dampened amygdala reactivity (76).

Our findings should be considered with limitations of the study in mind. First, the relatively small threat-only group limits our ability to make conclusions about the potential unique effects of threat experienced without co-occurring deprivation; however, this pattern is likely reflective of the broader population in which co-occurrence of threat and deprivation exposures are relatively common (7,77). We focused our neuroaffective analysis on a small number of regions of interest with prior support for ELA-related alterations; future work may consider other approaches that incorporate whole-brain analysis. Additionally, emerging evidence suggests that ELA may impact reward and threat *learning* specifically (78), pointing to the potential value of future work examining how ELA impacts affective learning processes during this developmental period.

In conclusion, we found that youth with experiences of deprivation or threat during childhood show delayed neurocognitive maturation compared to their socioeconomically matched peers. Isolated impacts of these experiences on neuroaffective processing should be further studied. Additionally, while ELA may confer increased risk for later negative outcomes, many individuals that experience ELA do not develop significant psychopathology (79,80). As healthy cognitive development is central to multiple domains of resilience (81), our work helps future studies aimed at understanding the developmental factors that promote resilience following exposure to deprivation or threat.

## Acknowledgements

OK was supported by the National Institute on Drug Abuse K01 DA059598. KLM was supported by National Institute on Drug Abuse K23 DA060996. The content is solely the responsibility of the authors and does not necessarily represent the official views of the National Institutes of Health.

## Data availability

The authors do not have permission to share data. The ABCD data repository grows and changes over time. Data used in the preparation of this article were obtained from the Adolescent Brain Cognitive Development® (ABCD) Study (https://abcdstudy.org) Release 4.0 (for ELA-variables) and Release 5.1. This is a multisite, longitudinal study designed to recruit more than 10,000 children age 9–10 and follow them over 10 years into early adulthood. The ABCD Study® is supported by the National Institutes of Health and additional federal partners under award numbers U01DA041048, U01DA050989, U01DA051016, U01DA041022, U01DA051018, U01DA051037, U01DA050987, U01DA041174, U01DA041106, U01DA041117, U01DA041028, U01DA041134, U01DA050988, U01DA051039, U01DA041156, U01DA041025, U01DA041120, U01DA051038, U01DA041148, U01DA041093, U01DA041089, U24DA041123, U24DA041147. A full list of supporters is available at https://abcdstudy.org/federal-partners.html. A listing of participating sites and a complete listing of the study investigators can be found at https://abcdstudy.org/consortium_members/. ABCD consortium investigators designed and implemented the study and/or provided data but did not necessarily participate in the analysis or writing of this report. This manuscript reflects the views of the authors and may not reflect the opinions or views of the NIH or ABCD consortium investigators.

## Code availability

The scripts used to generate the results, tables, and figures are shared at https://github.com/okardan/Deprv_Thrt_neurocog_neuroaff.

## Supplementary Materials

### S1. Demographic matching

Demographic matching was performed iteratively with the goal of creating matched groups for each exposure group drawn from the pool of participants with no reported deprivation or threat experiences. The matching was done for parental education, family income, age at Y0 and Y2, biological sex, and race/ethnicity of the participants. The iterations were to minimize the number of participants within exposure groups that had to be removed in order to achieve |Cohen’s ds| ≲ 0.1 between each exposure group and its final control group across all matching categories.

<u>Within the neurocognitive development analyses</u>, a total of 693 participants in the included sample reported exposure to deprivation only. To make the matching possible, 55 of these participants had to be excluded from the deprivation group (final N = 638). A total of 207 in the included sample reported exposure to threat only. Again, to make the matching possible 9 of these had to be excluded from the threat group (final N = 198). Finally, a total of 414 in the included sample reported exposure to threat non-exclusively (i.e., threat+), with 32 removed during matching (final N = 382). After one-to-one matching, duplicate participants in the matched control groups were removed, resulting in Ns = 534, 195, and 346 for the control groups of deprivation, threat, and threat+ groups, respectively (see Tables 1, 2, and 3).

<u>Within the neuroaffective development analyses</u>, 764 of the participants reported deprivation only. To make the matching possible with |Cohen’s ds| ≲ 0.1 between the groups across all matching categories, 43 of these had to be excluded from the deprivation group (final N = 721). A total of 239 participants reported threat exposure exclusively, 7 of which were removed during matching (final N = 232). A total of 494 participants reported threat or threat + deprivation, and 30 of these were removed to make the matching possible, resulting in 464 participants for the threat+ group. After one-to-one matching, duplicate participants in the matched control groups were removed, resulting in Ns = 603, 228, and 423 for the control groups of deprivation, threat, and threat+ groups, respectively (see Tables 1, 2, and 3).

Importantly, participants excluded during matching did not have higher intensity of exposure to deprivation or threat compared to the included participants (all ps > 0.25 in two-sample t-tests).

### S2. Adversity exposure variables

We utilized a multi-measure, multi-informant approach to calculating experiences of deprivation and threat that was informed by theory (e.g., (McLaughlin et al., 2021)) and prior topical work with the ABCD Study sample (e.g., (Breslin et al., 2025; Stinson et al., 2021)). Individuals with presence of one or more of these indicators at baseline, Year 1 follow-up, or Year 2 follow-up were classified as positive cases (i.e., deprivation exposure or threat exposure). Within deprivation and threat groupings, missing data were allowed for individuals with at least one positive indicator as their group assignment would remain the same regardless of additional data. However, because additional data could result in a change in group membership for youth in the unexposed group, youth were required to have data from all timepoints to be included in the matched control group.

#### Deprivation

Five youth-reported items from the Acceptance scale of the Children’s Report of Parental Behavior Inventory (Barber & Olsen, 1997; Schaefer, 1965) were used as a proxy measure of emotional deprivation. Youth reported on their parent/primary caregiver who was participating in the study on items such as “Makes me feel better after talking over my worries with them.” Options were “not at all like them,” “somewhat like them,” and “a lot like them.” In line with procedures described in (Stinson et al., 2021), a response of “not at all like them” on two or more of the five items was considered a positive case (potential deprivation). This measure was administered at baseline (ages 9-10) and Year 1 follow-up (ages 10-11).

Two youth-reported items from the Parental Monitoring Questionnaire (Karoly et al., 2016) were used as a proxy of supervisory deprivation based on their content and their frequency of use in prior adversity-related work in the ABCD Study sample (Breslin et al., 2025). These items measured parental knowledge of the youth’s whereabouts and youth’s ability to contact the parent when home without parent and were rated on a 5-point scale (1=“never” to 5=“almost always”). Items were dichotomized as detailed in (Stinson et al., 2021) with responses of 1 or 2 coded as potential deprivation (1) and responses of 3, 4, or 5 coded as no deprivation (0). This measure was administered at baseline, Year 1 follow-up, and Year 2 follow-up.

Four parent-reported items assessed experiences of adversity during the past 12 months due to financial hardship. Items ask about needing food but being unable to afford it, being evicted, having gas or electricity turned off, and needing to see a doctor but not going due to inability to afford it. We included these items as proxies of physical deprivation with a positive endorsement on one or more of the four items indicating a positive case. While these items are related to socioeconomic resources, they reflect direct experiences of deprivation that may result from limited socioeconomic resources. This measure was administered at baseline, Year 1 follow-up, and Year 2 follow-up.

#### Threat

Ten parent-reported items from the Kiddie Schedule for Affective Disorders and Schizophrenia for School-Aged Children (KSADS-COMP) assessed experiences of threat including items measuring verbal threats of death, physical abuse, sexual abuse, and witnessing domestic or community violence. Endorsement of one or more items was considered a positive case. This measure was administered at baseline and Year 2 follow-up.

### S3. Resting-state and task fMRI data

*<u>Resting-state fMRI data (rsfMRI)</u>* at Y0 and Y2 were used to construct a measure of cortical network maturation for the neurocognitive analyses (see variables section). In the ABCD Study, resting-state fMRI was acquired in four runs (∼5□min per run, full details are described in (Hagler et al., 2019)). The rsfMRI data were preprocessed using the pipeline described in (Sripada et al., 2021), which includes FreeSurfer normalization, ICA-AROMA denoising, CompCor correction, and censoring of high motion frames with a 0.5□mm framewise displacement threshold. Visual quality control (QC) was conducted to assess registration and normalization steps. To be included in the neurocognitive analyses, a participant needed to have at least one resting-state run at each timepoint (Y0 and Y2) with each run having more than 250 degrees of freedom left in the BOLD timeseries after confound regression and censoring. After preprocessing, rsfMRI data were spatially down-sampled to 333 parcels in the Gordon parcellation for the cortical regions (Gordon et al., 2016) and cortical functional connectivity matrices were generated for each participant for Y0 and Y2 by Fisher Z transformation of the 333*333 correlation matrices.

*<u>Monetary incentive delay (MID) task</u>* (Knutson et al., 2000) fMRI data were acquired in two runs (_∼_6 min per run; details are described in (Casey et al., 2018; Hagler et al., 2019)). Briefly, a trial of the MID task begins with an incentive cue (2000□ms) of five possible trial types (Win $.20, Win $5, Lose $.20, Lose $5, $0-neutral or no money at stake) and is followed by an anticipation event jittered to last between 1500 to 4000□ms. A target then appears for 150–500□ms, during which the participant responds to either win money or avoid losing money. The ABCD Data Analysis, Informatics & Resource Center (DAIRC) applied a general linear model to estimate individual subject level brain activity for contrasts between these trial types for brain regions of interest (ROIs). Cortical and subcortical ROIs were generated using the FreeSurfer brain imaging software package by DAIRC. We used the contrast beta weights for anticipation of large reward versus neutral trials of MID task in the Nucleus Accumbens and Caudate Freesurfer ASEG segmentation in both right and left hemispheres (ABCD 5.1 variable names: tfmri_ma_alrvn_b_scs_aalh and tfmri_ma_alrvn_b_scs_aarh, tfmri_ma_alrvn_b_scs_cdlh and tfmri_ma_alrvn_b_scs_cdrh). We excluded participants based on the DAIRC’s recommended task-specific exclusion criteria (variable imgincl_mid_include = 0), which indicates fewer than 200 degrees of freedom in the MID run, not passing their quality control, or unacceptable task performance.

*<u>Emotional n-back task (EN-back)</u>* task in the ABCD Study (Casey et al., 2018) includes 2 runs of 8 blocks each with 10 trials in each block. A picture is shown in every trial, and participants are tasked with indicating whether the picture is a “Match” or “No Match” to either 2 pictures prior (2-back) or a target picture shown at the beginning of the block (0-back). The task includes four blocks of each stimuli type (happy, fearful, and neutral facial expressions and places). Cortical and subcortical ROIs were generated using the FreeSurfer brain imaging software package by DAIRC. Negative emotional reactivity was estimated based on the contrast beta weights of fearful face vs. neutral face trials in the EN-back task in amygdala using Freesurfer segmentation and insula based on the Desikan-Killiany cortical atlas in both right and left hemispheres (ABCD 5.1 variable names: tfmri_nback_all_224, tfmri_nback_all_238, tfmri_nback_all_780, and tfmri_nback_all_814). We excluded participants based on the DAIRC’s recommended task-specific exclusion criteria (variable imgincl_nback_include = 0).

